# Haploinsufficiency of autism candidate gene NUAK1 impairs cortical development and behavior

**DOI:** 10.1101/262725

**Authors:** Virginie Courchet, Amanda J Roberts, Peggy Del Carmine, Tommy L. Lewis, Franck Polleux, Julien Courchet

**Author notes:** Co-corresponding authors Address correspondence to: Franck Polleux, Ph.D., Columbia University, Department of Neuroscience, Mortimer B. Zuckerman Mind Brain, Behavior Institute, Kavli Institute for Brain Science, 550 W 120^th^ Street, Room 1108 Northwest Corner Building, New York NY 10027, USA, Julien Courchet, DVM, Ph.D., Institut NeuroMyoGène, UMR5310 – U1217, Université Claude Bernard Lyon 1, Bâtiment Mendel 2e étage, 16 rue Raphael Dubois, F-69100 Villeurbanne, FRANCE.

## Abstract

Recently, numerous rare *de novo* mutations have been identified in children diagnosed with autism spectrum disorders (ASD). However, despite the predicted loss-of-function nature of some of these *de novo* mutations, the affected individuals are heterozygous carriers, which would suggest that most of these candidate genes are haploinsufficient and/or that these mutations lead to expression of dominant-negative forms of the protein. Here, we tested this hypothesis with the gene *Nuak1*, recently identified as a candidate ASD gene and that we previously identified for its role in the development of cortical connectivity. We report that *Nuak1* is happloinsufficient in mice in regard to its function in cortical axon branching *in vitro* and *in vivo. Nuak1*^+/−^ mice show a combination of abnormal behavioral traits ranging from defective memory consolidation in a spatial learning task, defects in social novelty (but not social preference) and abnormal sensorimotor gating and prepulse inhibition of the startle response. Overall, our results demonstrate that *Nuak1* haploinsufficiency leads to defects in the development of cortical connectivity and a complex array of behavorial deficits compatible with ASD, intellectual disability and schizophrenia.

---

Neurodevelopmental disorders form a group of disabilities resulting from impairment of brain development and include syndromes such as Autism Spectrum Disorders (ASD), intellectual disability, Attention Deficit/Hyperactivity Disorder (AD/HD) and schizophrenia. The etiology of neurodevelopmental disorders is complex as it involves a marked genetic contribution but probably also some environmental factors and/or a complex interaction between genes and environment ^1–4^. Despite strong heritability, less than 10% of syndromes such as ASD can be attributed to dominant mutations in a single gene ^1^. Recent studies revealed a large array of rare, *de novo* mutations in ASD patients ^5,6^. Yet their biological significance remains elusive largely because, despite the predicted loss-of-function nature of some of these *de novo* mutations, the affected individuals are heterozygous carriers, suggesting that most of these candidate genes are either haploinsufficient and/or that these mutations lead to expression of dominant-interfering forms.

The complex network of interconnected neurons forming the human brain results from the coordinated activation of precise cellular processes such as neurogenesis, cell migration, axon guidance, dendrite patterning and synaptogenesis, culminating in formation of functional neural circuits. Despite the important morphological and functional differences between neurons, axon development can be operatively divided in three stages ^7^: (1) axon specification, corresponding to the commitment of one neurite toward axonal fate, (2) axon elongation and guidance toward its target, and (3) branch formation, stabilization or elimination to form proper synaptic contacts. The cellular and molecular mechanisms underlying the establishment of neuronal connectivity have been extensively studied. Extracellular factors such as axon guidance cues, neurotrophins and other growth factors convey information provided by the local environment that is relayed to intracellular signaling molecules to coordinate axon morphogenesis through the triggering of local cellular processes. One such signaling relay, the serine/threonine kinase LKB1, is involved in several aspects of axonal development in the mouse cortex. *Lkb1* is a tumor suppressor gene encoding a “master kinase” that activates a group of 14 downstream effectors forming a specific branch of the kinome and related to the metabolic regulator AMPK ^8,9^. LKB1 exerts sequential functions in neuronal development through the activation of at least two distinct sets of AMPK-related kinases (AMPK-RK): neuronal polarization and axon specification through the kinases BRSK1/2 (SAD-A/B) ^10–12^, and axon elongation and terminal branching through NUAK1 (ARK5/OMPHK1) ^13^. Furthermore the BRSK ortholog SAD-1 is involved in synapse formation in *C. elegans* ^14,15^ whereas in the mouse cortex LKB1 has been shown to regulate synaptic transmission and neurotransmitter release through its ability to control mitochondrial Ca^2+^ uptake ^16^.

Several genes belonging to the AMPK-RK family came out of recent unbiased genetic screens for rare *de novo* mutations associated to neurodevelopmental disorders (Supplementary Table 1) ^17–19^. Yet the functional consequences of these mutations and to what extent they contribute to the physiopathology of the disease remain unknown. To address this question, we focused on *Nuak1*, one of the AMPK-RK genes strongly expressed in the developing brain ^13,20^. Mutations in the *Nuak1* genes have been linked to ASD ^17,21^, cognitive impairment ^19^ or AD/HD ^22^. Interestingly the patients carrying these rare *de novo* mutations are by definition heterozygous carriers. Therefore, the actual contribution of these mutations to the disease mechanisms suggests that *Nuak1* must be either haploinsufficient and/or the mutated allele must be exerting a dominant-negative influence over the wild-type allele.

In this study we present evidence demonstrating that the inactivation of one allele of the *Nuak1* gene results in a complex phenotype including growth retardation, defective axonal mitochondria trafficking and decreased terminal axon branching of cortico-cortical projections. We reveal that *Nuak1* heterozygosity disrupts long-term memory consolidation, alters the preference for social novelty without affecting conspecific social preference, and decreases the prepulse inhibition (PPI) of the acoustic startle reflex. Finally we adopted a gene-replacement strategy to further characterize the functional consequence of *Nuak1* rare *de novo* mutation identified in ASD patients. Overall, our results indicate that *Nuak1* is haploinsufficient for the development of cortical connectivity and support the hypothesis that mutation in *Nuak1* participates in the etiology of neurodevelopmental disorders.

## RESULTS

### NUAK1 haploinsufficiency impairs mouse cortical development

We first mapped the temporal and spatial pattern of expression of *Nuak1* in the developing and adult mouse brain using *in situ* hybridization. This revealed that *Nuak1* mRNA is highly expressed in the developing and adult mouse brain, with specific expression in the neocortex, pyriform cortex, hippocampus and cerebellum (granule cells but not Purkinje cells). In contrast, *Nuak1* expression was low or absent in the dorsal thalamus and striatum (Fig. S1A-I). Western-blot analyses from mouse brain extracts confirmed that NUAK1 protein is expressed at high level in the developing and adult cortex. In the hippocampus, NUAK1 expression was maintained throughout all developmental stages, although it decreased with age (Fig. S1J).

Constitutive knockout (KO) of *Nuak1* is lethal in the late embryonic/perinatal period in mice due to an abdominal midline closure defect called omphalocoele ^20^. Despite strong expression in the embryonic brain, the initial characterization of *Nuak1* KO embryos did not reveal any obvious morphological brain defect ^20^. Accordingly, we did not observe any difference between wild-type (WT), heterozygous (HET) and KO embryos in axonal markers (SMI312) or cortical lamination (TBR1: layer 6, and CTIP2: layer 5), indicating no major abnormalities in neuronal polarization, axon tract formation, neurogenesis, neuronal migration and lamination (Fig. S2A-F). We observed a marked decrease (~50%) of NUAK1 protein level in HET neurons (Fig. 1A), suggesting that the remaining allele does not compensate for the loss of one copy of *Nuak1*. Unlike the homozygotes constitutive knockout (NUAK1^−/−^), heterozygous mice (NUAK1^+/−^) are viable and fertile and do not display the omphalocoele phenotype. Quantitative analysis revealed a moderate but significant growth retardation in NUAK1^+/−^ compared to WT littermates (Fig. 1B-C), more pronounced in males than females.

**Figure 1:**
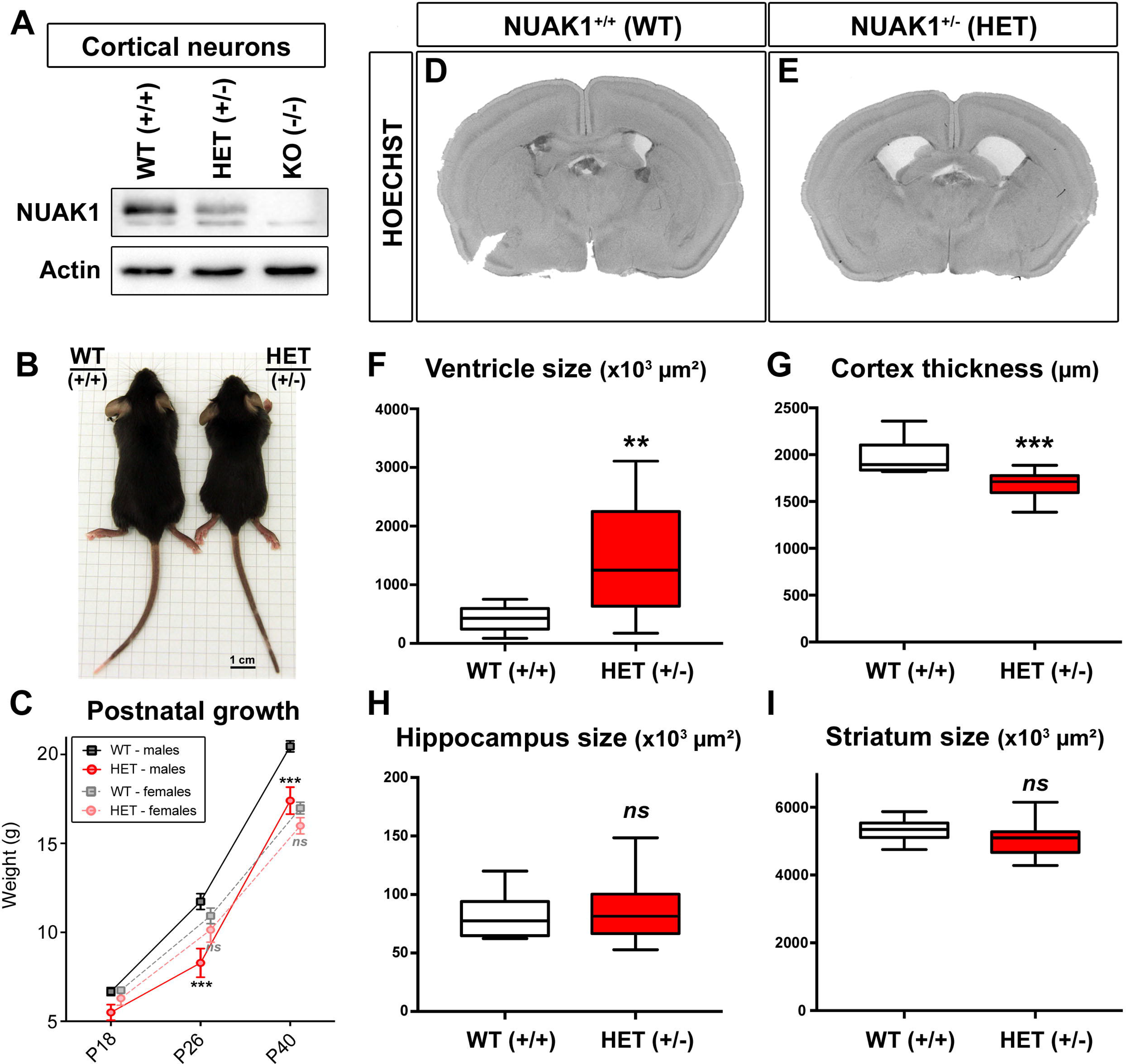
Growth retardation and hydrocephaly in NUAK1 mutant mice. (A) Western-blot analysis of NUAK1 expression in 5DIV cortical neurons of WT, HET and KO animals. (B) Body size difference between males WT and HET NUAK1 mice at age P26. (c) Postnatal body weight growth curve of WT and HET NUAK1 mice before (P18), or after (P26-P40) weaning. Average ± SEM. N_WT-females_=17, N_HET-females_=14, N_WT-males_=14, N_HET-males_=8. Analysis: 2way ANOVA with Bonferroni’s multiple comparisons. (D-E) Representative coronal sections of P21 NUAK1 WT (D) and HET (E) mice stained with Hoechst dye. (F-I) Measurement of ventricle size (F), cortex thickness (G), hippocampus (H) and striatum size (I) from P21 coronal sections of NUAK1 WT and HET animals. Data represents median, 25^th^ and 75^th^ percentile. Analysis: Two-tailed unpaired T-test. (F, G and I) N_WT_=10, N^HET^=15. (H) N_WT_=7, N^HET^=13.

We performed histochemical analyses of NUAK1^+/−^ mouse brains at Postnatal day (P)21 (weaning) or P40 (young adult). The global organization of the brain appeared largely normal, with a normal segregation of somatodendritic and axonal markers (Supplementary Fig. 2G-H) and no difference in the distribution of neuronal layers (Supplementary Fig. 2I-L) between WT and HET animals. Yet the most striking observation was a marked enlargement of the lateral ventricles in NUAK1^+/−^ mice never observed in WT littermates on this C57Bl/6J genetic background (Fig. 1D-E and Fig. S2G-H). The size of lateral ventricles was nearly doubled in NUAK1^+/−^ mice (Fig. 1F), which was accompanied by a corresponding thinning of the cortex along its radial axis (Fig. 1G), whereas the volume of two other brain structures (the striatum and hippocampus) was not affected (Fig. 1H-I). Overall, our data suggest that although NUAK1^+/−^ mice are viable, *Nuak1* is haploinsufficient regarding brain development.

### NUAK1 exerts a dose-dependent role in cortical axon development

We previously characterized that NUAK1 is necessary and sufficient for terminal axon branching of layer 2/3 cortical neurons through the presynaptic capture of mitochondria ^13^. In order to determine if these phenotypes are also observed upon loss of one copy of *Nuak1*, we first performed neuronal cultures following *ex utero* cortical electroporation (EUCE) of WT, NUAK1^+/−^ and NUAK1^−/−^ embryos at E15.5. We observed a reduction of axon length and branching in NUAK1^+/−^ and NUAK1^−/−^ neurons when compared to WT neurons (Fig. 2A-C). Time-lapse imaging of mitochondria trafficking in developing layer 2/3 cortical axons revealed that mitochondria motility is markedly increased in the anterograde and retrograde direction for both NUAK1^+/−^ and NUAK1^−/−^ neurons compared to control WT neurons (Fig. 2D-F). Quantifications confirmed that NUAK1 exerts a dose-dependent effect on axon development and that HET neurons have a significant reduction in both maximum axon length and the number of collateral branches (Fig. 2G-I). We also quantified the fraction of stationary mitochondria and observed a marked reduction in the percentage of stationary mitochondria per axon in NUAK1^+/−^ and NUAK1^−/−^ neurons compared to control WT neurons (Fig. 2J).

**Figure 2:**
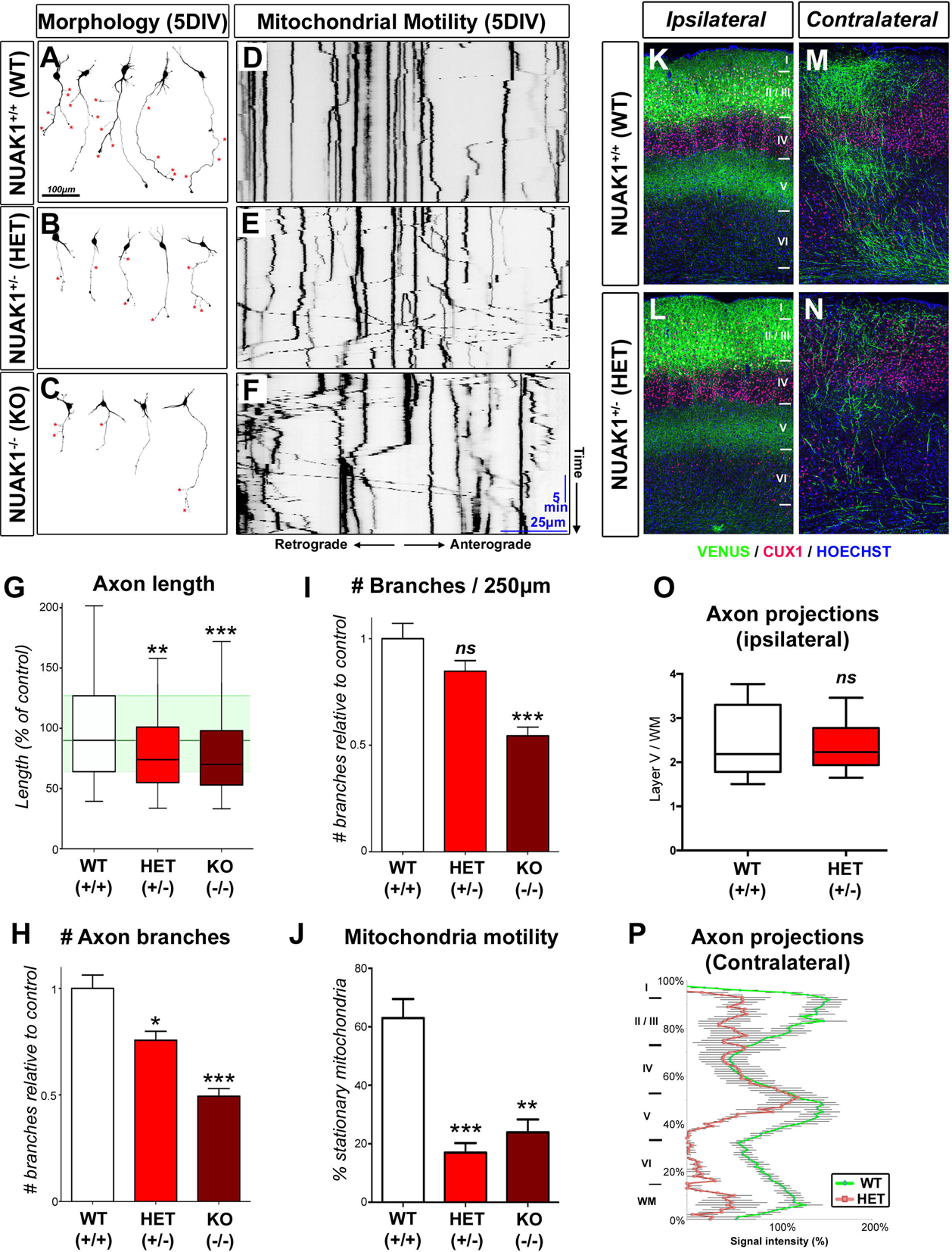
NUAK1 is haploinsufficient for axon development. (A-C) Representative neurons imaged after 5DIV in WT, HET and KO animals. Neuron morphology was visualized through mVenus expression. (D-F) Representative kymographs of axons in WT, HET and KO neurons at 5DIV. Mitochondria were visualized through expression of mito-DsRed. (G-I) Quantifications showed a gradual reduction in axon length (G) and collateral branches (H-I) in HET and KO animals. (G) Data represents median, 25^th^ and 75^th^ percentile. (H-J) Average ± SEM. N_WT_=127, N_HET_=232, N_KO_=203. Analysis: one way ANOVA with Bonferroni’s post test. (J) Quantification of mitochondria motility in the axon. Average ± SEM. N_WT_=12, N_HET_=9, N_KO_= 13. Analysis: Kruskal-Wallis test with Dunn’s multiple comparison. (K-N) Higher magnification of the ipsilateral (K-L) or contralateral side (M-N) of P21 WT or HET mice following in utero cortical electroporation of mVenus. Blue: Hoescht, Red: CUX1 immunostaining. (O-P) Quantification of normalized mVenus fluorescence in Layer 5 of the ipsilateral cortex (median, 25^th^ and 75^th^ percentile) (O) and along a radial axis in the contralateral cortex (Average ± SEM) (P). N_WT_=6, N_HET_=5. Analysis: two tailed unpaired T-test (O).

We subsequently turned to long-term *in utero* cortical electroporation (IUCE) to determine the consequences of *Nuak1* heterozygosity on terminal axon branching *in vivo*. We targeted pyramidal neurons of the superficial layers 2/3 of the primary somatosensory cortex (S1) through IUCE of a mVenus coding plasmid at E15.5 in WT (NUAK1^+/+^) and HET (NUAK1^+/−^) embryos. Mice were sacrificed at Postnatal day (P)21, a stage when terminal branching of these cortico-cortical projecting (callosal) axons is adult-like ^23,24^. The loss of one copy of *Nuak1* had no effect on neurogenesis, radial migration and axon formation (Fig. 2K-L). Furthermore, callosal axons crossed the midline and reached the contralateral hemisphere. However we observed a marked decrease in terminal branching on the contralateral hemisphere in NUAK1^+/−^ mice compared to WT littermates (Fig. 2M-N). Interestingly, axon branching on the ipsilateral hemisphere was largely unaffected (Fig. 2O-P). A similar phenotype was observed in older mice (P90) mice (Fig. S3), indicating that this deficit in terminal axon branching does not result from a simple delay in axon development and persists in adulthood.

### NUAK1 heterozygosity disrupts long-term memory consolidation

Despite being involved in cortical axon morphogenesis (present study and ^13^), the functional consequences of *Nuak1* heterozygosity are largely unknown. In order to investigate the effect of NUAK1 haploinsufficiency on mouse behavior, we generated age-matched cohorts of WT and NUAK1^+/−^ mice and performed a battery of behavioral assays relevant to neurodevelopmental disorders. Control experiments demonstrated that vision and olfaction were not impaired in NUAK1^+/−^ mice, as demonstrated by tracking of head movement using an optomotor test (Fig. S4A) and by the habituation/dishabituation to novel odor test (Fig. S4B). Furthermore NUAK1^+/−^ mice exhibited no difference in spontaneous locomotor activity and exploratory behavior in the Open Field (Fig. S5A-C).

Next, we tested how NUAK1 heterozygosity affects memory formation and consolidation through two distinct cognitive paradigms: first, we investigated non-spatial memory using the Novel Object Recognition (NOR) task ^25^ (Fig. 3A). Adult NUAK1^+/−^ and WT littermates behaved similarly during the initial training period (data not shown). There were no significant sex or genotype effects in the subsequent test trial; however, there were effects of object contacts (F(1,35)=14.21, p<0.001) and contact times (F(1,35)=15.63, p<0.001), with both WT (p<0.005) and NUAK1^+/−^ (p<0.01) mice, showing increased interest in the novel object (Fig. 3B-C). Second, we tested spatial memory formation and maintenance using the Barnes Maze assay ^26^ (Fig. 3C). While this test is similar to the Morris Water Maze in probing spatial memory, it is less physically taxing and does not induce stress/anxiety response linked to swimming in water ^27^. There were no significant differences involving sex or genotype in the acquisition of escape behavior. In the probe trial both WT and NUAK1^+/−^ mice showed a clear preference for the target quadrant relative to other quadrants (Fig. 3D, probe test #1 [F(1,44)=11.1, p<0.005]). However, when retested in the same conditions after one month, while there was a significant effect of quadrant time (F(1,44)=6.9, p<0.01), it was only the WT mice that showed a significant preference for the target quadrant (p=0.005), while NUAK1^+/−^ mice did not (p=0.45; Fig. 3E, probe test #2). Altogether, these results indicate that NUAK1 partial loss-of-function disrupts long-term memory maintenance/consolidation.

**Figure 3:**
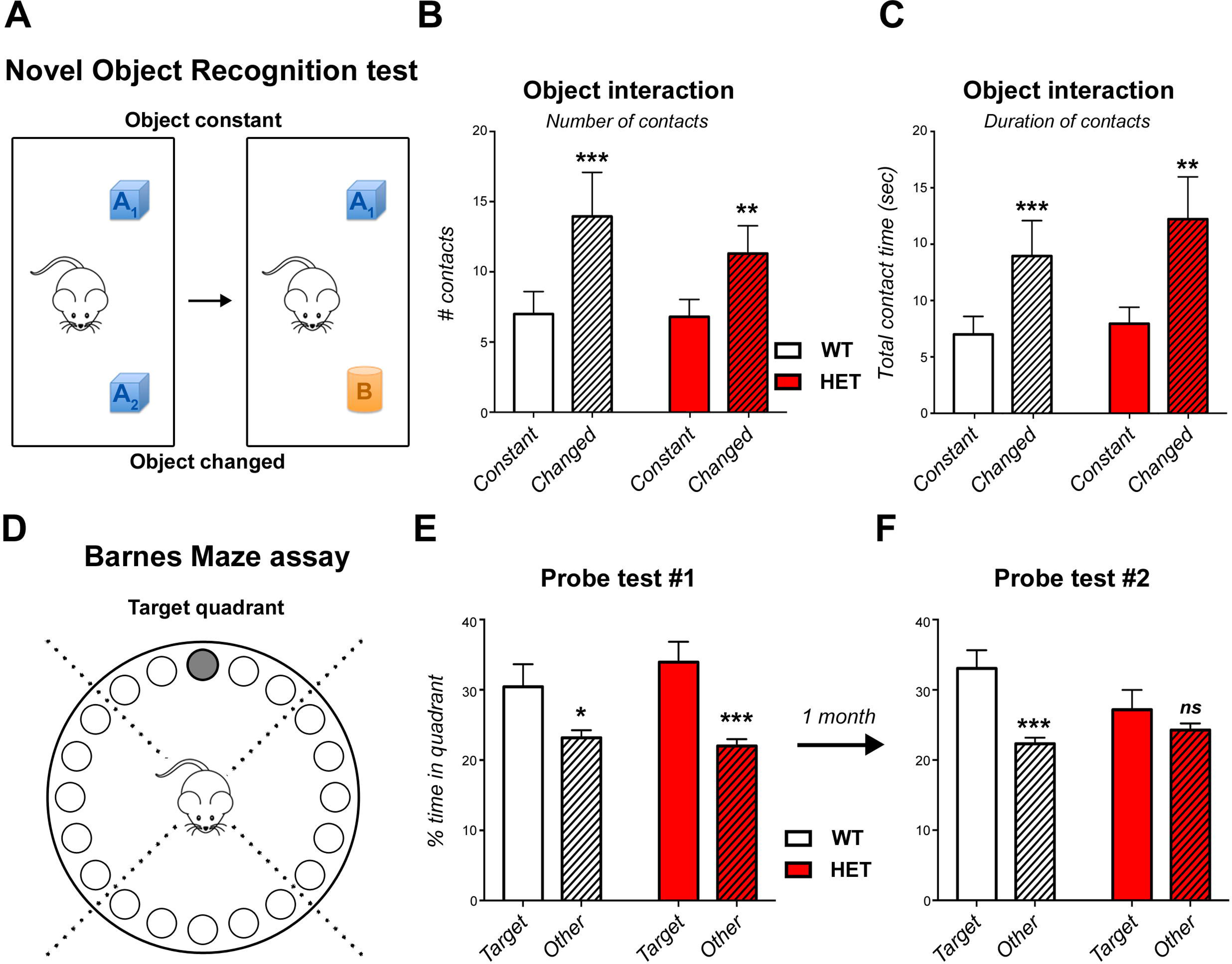
Long-term memory consolidation phenotype of NUAK1 HET mice. (A) Novel object assay design. Mice were exposed to two similar objects, followed by a change in one of the two objects. (B-C) Quantification of the number (B) and duration (C) of contacts during the novel object recognition assay. Average ± SEM. N_WT_=19, N_HET_=20. Analysis: 2-way ANOVA with Bonferroni’s multiple comparisons. (D) Barnes Maze spatial memory assay. (E) Quantification of time spent in target quadrant versus in other quadrants. (F) Mice were tested after 1 month to assess memory consolidation. Average ± SEM. Analysis: 2-way ANOVA with Bonferroni’s multiple comparisons.

### NUAK1 heterozygosity impairs novelty preference and sensory gating

We next performed a series of behavioral assays relevant to autistic-like traits. We observed spontaneous activity through Open Field sessions (Fig. S5A-F) but did not detect any differences between WT and NUAK1^+/−^ mice in behaviors classically associated with anxiety or repetitive behavior.

Social interactions were assessed using the 3-chamber sociability assay (Fig. 4A) ^28^. There was no overall genotype difference, but a significant overall effect of time in chamber (F(2,132)=227.6, p<0.0001) and direct contact times (F(1,66)=51.3, p<0.0001). NUAK1^+/−^ mice displayed the same preference, measured by the time spent in each chamber (Fig. 4B) and direct object/mouse contact time (Fig. 4C) as their WT littermates when offered a choice between another mouse and an inanimate object, indicating that conspecific sociability is not affected by NUAK1 heterozygosity. In a second test, we assessed the preference of NUAK1^+/−^ mice for social novelty (Fig. 4D). In this case, there were overall effects of chamber and contact times, but also significant interactions between these factors and genotype (F’s – 3.7−5.1, p<0.05). WT mice spent more time in the chamber with the novel mouse and interacted more with the novel individual. However, NUAK1^+/−^ mice displayed no preference between the familiar and the novel mouse (Fig. 4E-F). Importantly this effect was not due to a lack of interest for novelty since NUAK1^+/−^ mice showed a preference for a novel object similar to their WT littermates in the Novel Object Recognition (NOR) task ^25^ (Fig. 3A-C). Altogether, our results point to a deficit in the preference for social novelty upon loss of one copy of *Nuak1*.

**Figure 4:**
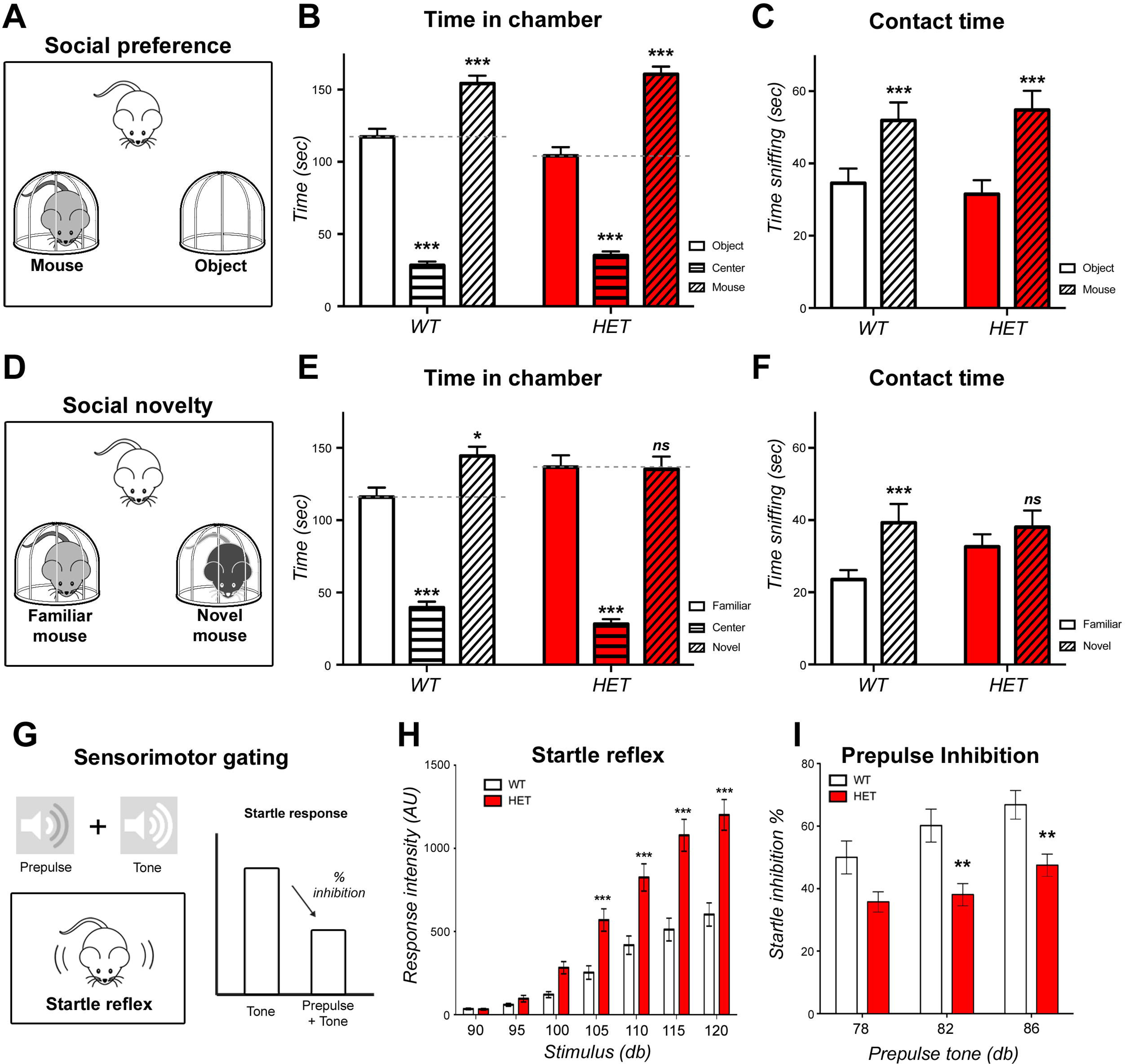
Sociability and sensorimotor phenotype of NUAK1 HET mice. (A) 3-chamber sociability assay design. (B-C) Quantification of time spent in each chamber (B) and contact time with the mouse or the empty cage (C) did not reveal any social deficit in NUAK1 HET mice. Data: Average ± SEM. N_WT_=36, N_HET_=34. Analysis: 2-way ANOVA with Bonferroni’s multiple comparisons. (D) Social novelty assay design. (E-F) Quantification of time spent in chambers (E) and contact time with a familiar and a novel mouse (F). Data: Average ± SEM. N_WT_=36, N_HET_= 34. Analysis: 2-way ANOVA with Bonferroni’s multiple comparisons. (G) Sensorimotor gating assay and measurement of the startle response. The startle reflex intensity (H) and % inhibition of the startle response upon pre-stimulation with a milder tone (I) showed altered response in NUAK1 HET. Average ± SEM. N_WT_= 24, N_HET_=24. Analysis: 2-way ANOVA with Bonferroni’s multiple comparisons.

We subsequently investigated sensorimotor gating through prepulse inhibition (PPI) of the startle response induced by weaker prestimuli ^29,30^ (Fig. 4G). We observed a marked enhancement of the startle response in NUAK1^+/−^ mice ([F(1,44)=24.9, p<0.0001], Fig. 4H). Interestingly, there was a significant startle sound level x genotype x sex interaction (F(7,308)=2.09, p<0.05), revealing a more pronounced increase in NUAK1+/− males compared to NUAK1+/− females (Fig. S6B-C).This increased startle response was accompanied by a significant reduction of the PPI ([F(1,44)=11.5, p=0.002], Fig. 4I), with stronger effect in males than in females (Fig. S6B-C). We retested the mice by submitting NUAK1^+/−^ to a milder tone in order to normalize the intensity of the startle response between WT and NUAK1^+/−^ animals. Under these conditions, we observed a milder, and nonsignificant PPI reduction in NUAK1^+/−^ males and no differences in NUAK1^+/−^ females compared to control WT littermates (Fig. S6D-E, see figure legend for statistical results). Finally, we observed that the optimal InterStimulus Interval (ISI) was similar between WT and NUAK1^+/−^ mice, and that the PPI was impaired at all ISI examined, indicating that the deficit in PPI does not result from an abnormality in the temporal function of the startle reflex (Fig. S6A). Taken together, our results indicate that NUAK1 heterozygosity induces a deficit in the PPI reflex that is likely due to a strong increase in the initial startle response.

### NUAK1 cell-autonomously control axon branching

Given the embryonic lethality characterizing the NUAK1^−/−^ constitutive knockout, we generated a conditional floxed *Nuak1* mouse model (NUAK1^F^ allele) (Fig. S7A-B) in order to compare partial and complete genetic loss of *Nuak1* function during development of cortical connectivity at postnatal stages. We first performed primary cortical neuron cultures coupled with EUCE of plasmids encoding Cre recombinase. The loss of one (NUAK1^F/+^) or two (NUAK1^F/F^) allele of *Nuak1* showed again a dose-dependent effect on axon growth and branching (Fig. S7C-H), thus validating the conditional allele as a potent loss-of-function mutation phenocopying the constitutive allele.

Next, we performed long-term IUCE of plasmids encoding Cre recombinase in floxed NUAK1 embryos at E15.5 to quantify axon branching of layer 2/3 pyramidal neurons *in vivo* at P21. Importantly, this approach is also testing the cell-autonomy of NUAK1 deletion since Cre-expressing neurons are relatively sparse (~10% of all layer 2/3 neurons) and their axons grow in a largely wild-type environment. Similarly to our observations in NUAK1^+/−^ mice, *Nuak1* conditional deletion had no effect on neurogenesis, radial migration, axon formation and ipsilateral branching (Fig. 5A-C and 5G). However, we observed a marked decrease of contralateral axon branching of layer 2/3 neurons upon inactivation of one or both alleles (Fig. 5D-F and 5H), strongly arguing that *Nuak1* is haploinsufficient with regard to the establishment of cortico-cortical connectivity during postnatal development.

**Figure 5:**
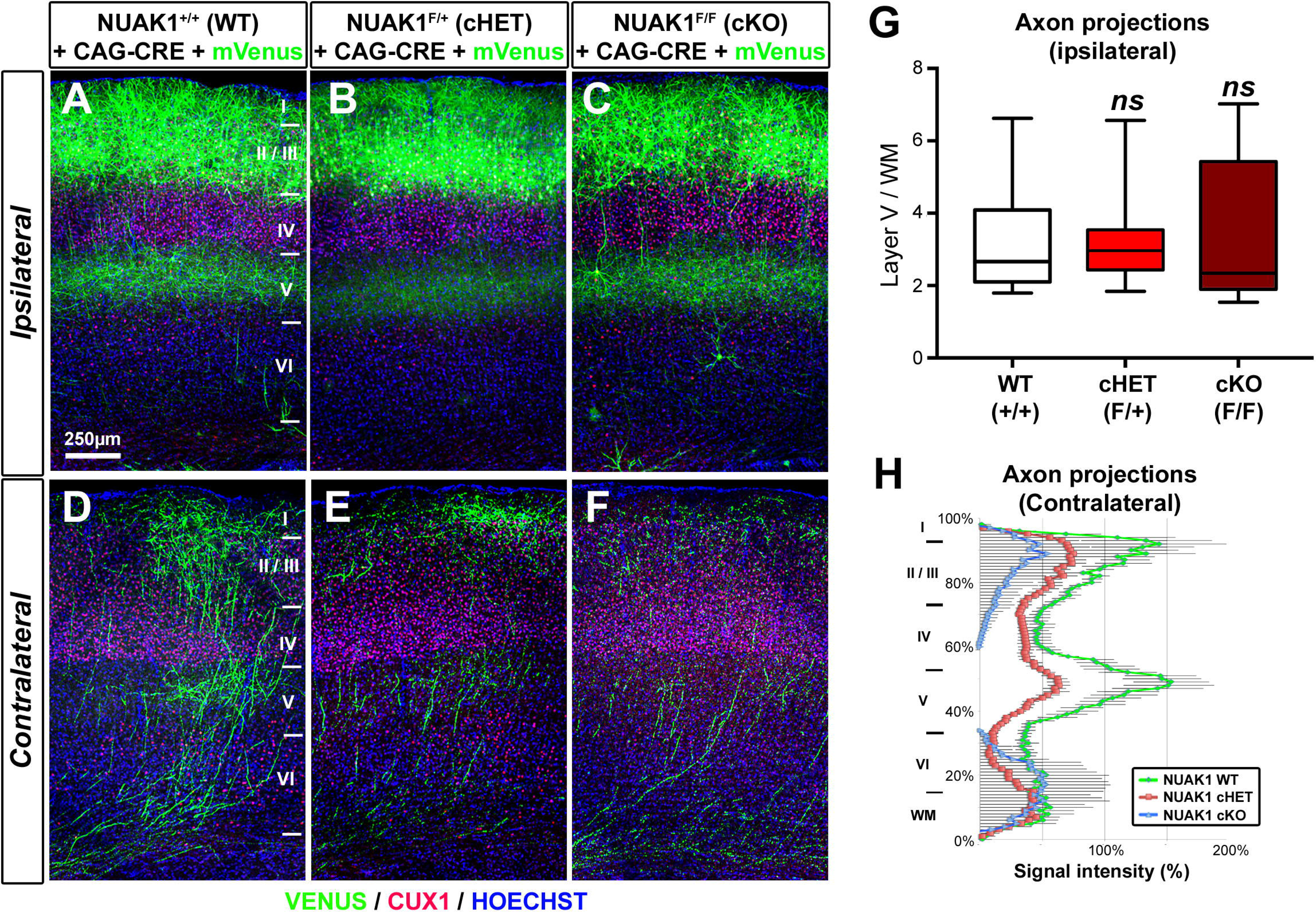
Cell autonomous reduction in axon branching of NUAK1-deficient neurons. (A-F) Histochemistry of the ipsilateral or contralateral side of NUAK1^+/+^, NUAK1^F/+^ and NUAK1^F/F^ mice at P21 following in utero electroporation with CRE and the fluorescent protein mVenus. Blue: Hoescht, Red: CUX1 immunostaining. (G-H) Quantification of normalized mVenus fluorescence in Layer 5 of the ipsilateral cortex (G) and along a radial axis in the contralateral cortex (H). Data: Average ± SEM. N_WT_=6, N_HET_=13, N_KO_= 6. Analysis: Mann-Whitney test.

Finally in order to determine the consequence of a complete deletion of NUAK1 on mouse behavior, we crossed NUAK1^F/F^ mice with NEX^CRE^ mice to drive CRE recombinase expression specifically in neural progenitors of the dorsal telencephalon^31^ (Fig. 6A). Conditional inactivation of one (cHET) or both (cKO) copies of *Nuak1* had no effect on mouse viability and postnatal growth (Fig. 6A and S8A). We then performed the NOR task and the three-chambered social preference and social novelty assay on age matched cohorts of WT, cHET and cKO animals. As was the case upon constitutive deletion of NUAK1, there was no overall effect of genotype on novel object recognition (Fig. S8B) and conspecific sociability as measured by time in chamber (Fig. 6B) and direct contact times (Fig. S8C). However during the social novelty phase of the assay, NUAK1 cHET and NUAK1 cKO mice showed a lack of preference for the novel mouse compared to the familiar mouse as measured by time in chamber (Fig. 6C) and direct contact times (Fig. S8D). Taken together our results indicate that the partial or complete inactivation of *Nuak1* in the dorsal telencephalon disrupts the preference for social novelty in the mouse and confirms that NUAK1 is haploinsufficient for mouse social behavior.

**Figure 6:**
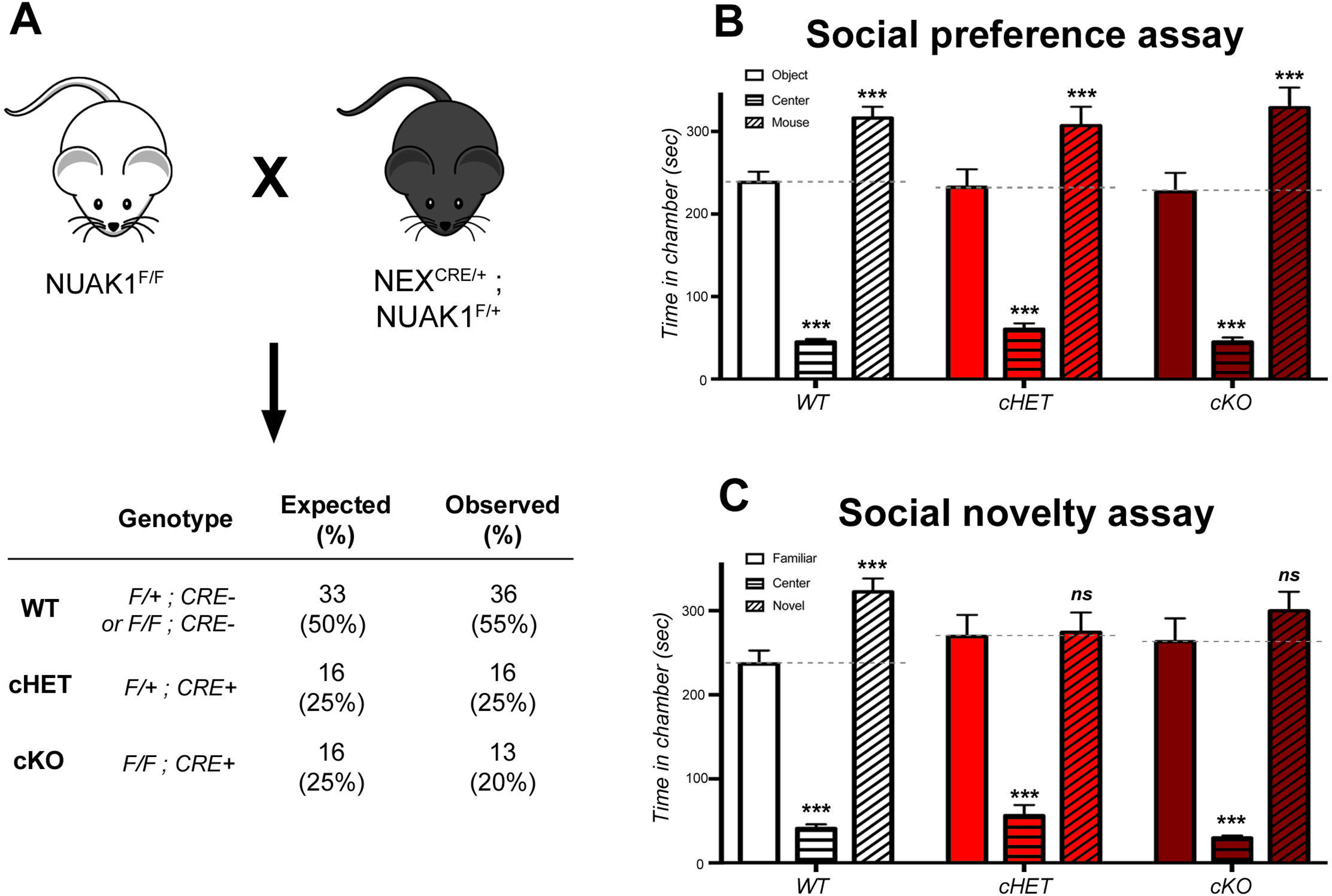
Reduced preference for social novelty upon complete loss of NUAK1 in the cortex. (A) Breeding strategy for dorsal-telencephalon specific inactivation of *Nuak1*. Observed proportion of each genotype was matching the expected proportion following mendelian rule. (B-C) 3-chamber sociability and social novelty assay in NUAK1 cKO mice. Quantification of time spent in each chamber revealed a decreased preference for social novelty in cHET and cKO mice. Data: Average ± SEM. N_WT_=24, N_cHET_= 8, N_cKO_=9. Analysis: 2-way ANOVA with Bonferroni’s multiple comparisons.

### Functional analysis of NUAK1 mutation associated with ASD

Finally, we tested whether NUAK1 mutations identified in autistic patients affect the function of the protein. The *de novo* mutation most frequently identified in genetic studies and with the highest confidence score is a nonsense mutation inducing a Premature Termination Codon (PTC) corresponding to Glutamine 433 (mouse 434) of the protein (Fig. 7A and Supplementary Table 1). Notably this PTC occurs in the long, last coding exon of *Nuak1* (Fig. S7A) and thus is not predicted to be subjected to nonsense mediated decay ^32^. The mutation (hereafter named NUAK1 STOP) predictably leads to the truncation of a large portion of the C-terminal extremity (CTD) of the protein, and removes one activating AKT-phosphorylation site as well as two GILK motifs required for the interaction with the myosin phosphatase complex ^33^.

**Figure 7:**
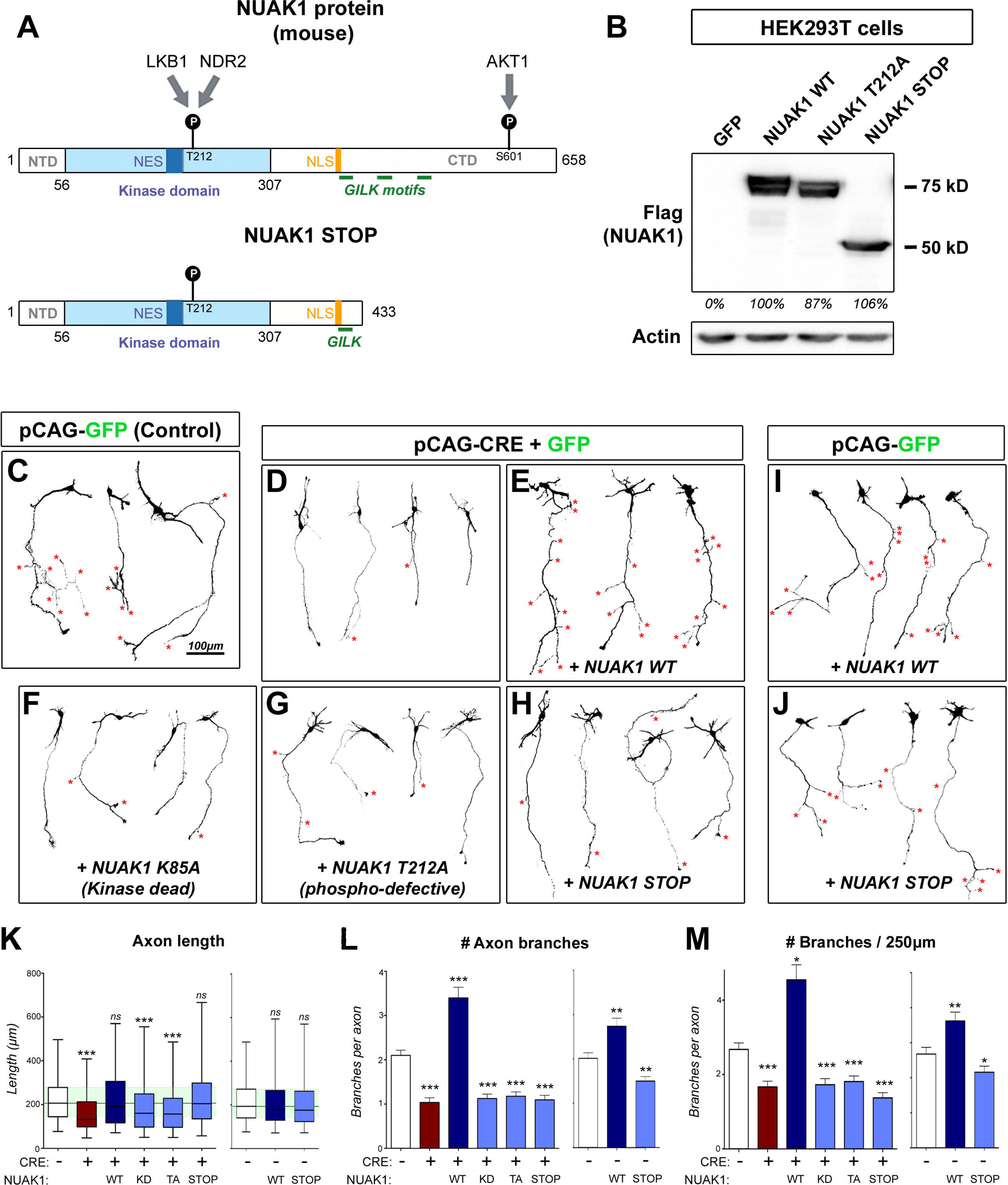
Functional analysis of NUAK1 mutations through a protein-replacement strategy. (A) Structure of the NUAK1 protein (mouse) and predicted structure of the Q434STOP mutant. Known or predicted domains are indicated. NTD: N-terminal domain. CTD: C-terminal domain. NES: Nuclear Export Sequence (predicted). NLS: Nuclear Localisation Sequence (predicted). (B) Validation of NUAK1 mutant expression in HEK293T cells by Western-blot. (C-J) Representative neurons imaged after 5DIV following ex vivo electroporation of the indicated plasmids in NUAK1^F/F^ animals. Neuron morphology was visualized through mVenus expression. (K-M) Quantification of axon length (K), number of branches per axon (L) and normalized branch number (M) in neurons from (C-J). Quantifications were normalized to the average value of Control (CRL) condition to allow comparison of distinct, independent experiments. Data represents median, 25^th^ and 75^th^ percentile (K) or Average ± SEM (L-M). N_CTL_=189, N_CRE_=138, N_CRE_+_WT_=90, N_CRE_+_KD_=150, N_CRE_+_TA_=174, N_CRE_+_STOP_=125, N_WT_=134, N_STOP_= 198. Analysis: Kruskal-Wallis test with Dunn’s multiple comparison.

We used site-directed mutagenesis to introduce the PTC mutation in NUAK1 coding plasmids, creating an expression vector encoding NUAK1 STOP, and used this construct to determine the consequences of NUAK1 protein truncation. We observed by Western-blot that the mutation had no impact on the expression of the protein (Fig 7B), suggesting that the removal of a large portion of NUAK1 CTD is unlikely to reduce protein stability.

We then investigated the consequences of expression of NUAK1 STOP protein on axon growth and branching. To reach that goal, we adopted a gene-replacement strategy to express either WT or mutant NUAK1 STOP protein in NUAK1-null neurons. We performed EUCE in NUAK1^F/F^ embryos at E15.5 and determined axon morphology at 5DIV. As reported above upon the inactivation of NUAK1 expression through Cre recombinase expression, neurons had shorter axons with reduced branching (Fig. 7C-D). Re-expression of WT NUAK1 protein fully rescued axon length and axon branching (Fig. 7E). On the contrary two mutants, a kinase dead (KD) mutant that totally abolishes NUAK1 catalytic activity, and a mutant of the LKB1-activation site (TA), failed to rescue the axonal phenotype (Fig. 7F-G) confirming that NUAK1 function in axon morphogenesis is kinase-dependent. Interestingly, NUAK1 STOP mutant could rescue axon elongation to the same extent as the WT NUAK1, but was not able to rescue collateral branch formation (Fig. 7H and L-M).

Finally we investigated whether the NUAK1 STOP mutation acts as a dominant negative (DN). We expressed either WT or mutant forms of NUAK1 into WT neurons (NUAK1^F/F^ without Cre) by EUCE and observed axon morphology at 5DIV. As observed previously ^13^, overexpression of WT NUAK1 was sufficient to increase axon branching (Fig. 7I). In contrast, neurons expressing the NUAK1 STOP mutant protein had limited to no effect on axon length or branching (Fig. 7J). Quantification of axon morphology confirmed that NUAK1 STOP mutant failed to increase branching (Fig. 7K-M). We only observed a moderate reduction in collateral branch formation upon expression of the NUAK1 STOP mutant, indicating that NUAK1 truncation has only a moderate inhibitory activity over endogenous NUAK1 and behaves mostly as a loss-of-function.

## DISCUSSION

In the present study, we characterized for the first time the consequences of *Nuak1* heterozygosity on brain development and cortical circuits formation. Previous observations reported that *Nuak1* is highly enriched in the developing brain ^20,34^, yet its function in brain development was previously concealed by the embryonic lethality of constitutive *Nuak1* knockout mice. We now report that *Nuak1* expression remains elevated in the mouse brain at various stages of postnatal development. Furthermore our results demonstrate that the inactivation of one copy of *Nuak1* induces a significant cognitive and behavioral phenotype in mice.

The genetic architecture of neurodevelopmental disorders such as autism spectrum disorders (ASD) is emerging as complex and in the vast majority of cases seems to deviate from mono-allelic, recessive mutations. Recent evidence suggest that more often, rare *de novo* mutations have been identified in ASD patients including potential loss-of-function mutations in AMPK-RK such as *Nuak1* ^17,19,21^, *Sik1* ^21^, *Brsk1* ^18^ or *Brsk2* ^19^ However, functional assessment of the phenotypic impact of these rare *de novo* mutations is currently unknown. The present study demonstrates for the first time that *Nuak1* is happloinsufficient in regard to brain development implying most likely that patients carrying heterozygote loss-of-function mutations in this gene could cause defects in brain connectivity and a range of behavioral defects compatible with ASD, intellectual disability and schizophrenia.

Many of the AMPK-RKs are expressed in the developing brain and several have been linked to neurodevelopmental disorders. For example, mutations in *Sik1* have been identified in patients suffering from developmental epilepsy ^35^, and polymorphisms in *Mark1* are associated with an increased relative risk for ASD ^36^. AMPK-RKs might have partially overlapping function protecting the developing brain from deleterious and potentially lethal mutations. As an example the simultaneous inactivation of the polarity kinases BRSK1 and BRSK2 is required to disrupt axon specification in mouse cortical neurons yet *Brsk1;Brsk2* double KO is lethal shortly after birth ^12^ whereas single gene inactivation is less phenotypic and viable. We previously observed that NUAK1 function in axon outgrowth and branching is not shared by other AMPK-RKs such as AMPK, BRSK1/2 or NUAK2 ^13^. This lack of redundancy between NUAK1 and other AMPK-RKs in axon morphogenesis might explain why mutations in *Nuak1* could have important consequences on brain development.

LKB1 is the upstream activator of all 14 AMPK-RKs ^8^ and its activity is necessary for proper axon morphogenesis and brain development ^10,13^. Dominant mutations in *Lkb1* cause the genetically inherited Peutz-Jeghers syndrome (PJS) ^37^, a rare disease associating hamartomas of the gastrointestinal tract and increased risk of cancer. Yet unlike other hamartomatous polyposis syndromes such as Tuberous Sclerosis (Tsc1/2) or the PTEN Hamartoma Tumor Syndromes ^38,39^, the clinical manifestations of PJS do not typically include neurological manifestations ^37^. As it is the case for many tumor suppressor genes for which a two hit inactivation of genes is required for cell transformation, it is worth noting that LKB1 is expressed to almost normal levels in LKB1^+/−^ mice contrary to NUAK1 protein which is reduced by roughly 50% in NUAK1^+/−^ constitutive knockout mice. Catalytic activity and subcellular localization of LKB1 is dependent upon the formation of a heterotrimeric complex with the pseudokinase STRAD and the adaptor protein Mo25 ^40,41^. Interestingly genetic inactivation of STRADα (also called LYK5), one of the two isoforms of the STRAD pseudokinase, cause a rare neurodevelopmental syndrome termed PMSE (Polyhydramnios, Megalencephaly and Symptomatic Epilepsy) ^42,43^. Furthermore STRADα plays a dominant but partially redundant role over STRADβ in axon development ^44^. Therefore *Lkb1* might be indirectly linked to disorders of neurodevelopment through subtle alterations of the function of the LKB1 complex.

In this study, we adopted a gene-replacement strategy to determine the functional relevance of one of the *de novo* mutations in the *Nuak1* gene potentially to ASD in human and leading to the expression of a truncated protein. The expression of abnormal, truncated or mutant protein raises the question whether the observed biological effects result from a loss of a normal function, or a gain in a aberrant function of the protein. Our data suggest that the NUAK1 STOP mutation causes a loss of function form of the protein; indeed we did not observe any marked dominant-negative effect upon NUAK1 STOP over-expression in WT cortical neurons, which would be the expected outcome from a gain of an aberrant function. Interestingly the NUAK1 protein has a long C-terminal tail containing potential regulatory sites including an AKT phosphorylation site ^45^ and three short GILK motifs promoting NUAK1 interaction with Protein Phosphatase 1β (PP1β) ^33^. LKB1 activation is achieved through nuclear export and cytosolic capture within the STRAD/Mo25 complex ^40,41^. Similarly it is plausible that NUAK1 activity and subcellular localization depend upon protein-protein interactions through the C-terminal domain of the protein.

The behavioral alterations in NUAK1^+/−^ mice are largely consistent with the identification of rare *de novo Nuak1* mutations in autistic patients ^17,21^. Indeed NUAK1^+/−^ mice display some features associated with the autistic clinical spectrum, including disruption of social novelty preference, moderate cognitive deficits, and a reduction in the PPI reflex. However, other behavioral alterations typical of ASD were absent in NUAK1^+/−^ mice. In particular, we did not detect any stereotyped or repetitive behavior in heterozygous mice, nor an effect on general sociability. In humans, the large heterogeneity in the clinical manifestations of syndromes falling in the autistic spectrum partially overlaps clinical manifestations associated with psychiatric disorders such as schizophrenia ^46,47^. Importantly most behavioral alterations we report here are equally indicative of ASD, intellectual disability and schizophrenia-like disease. Of note the activity of NUAK1^+/−^ mice was largely normal despite reports linking *Nuak1* mutations and AD/HD ^22^. One important distinction between ASD and schizophrenia is the age of the onset of phenotype, schizophrenia being typically diagnosed at adolescence or in young adults following a rather asymptomatic prodromal phase, whereas autism is typically detected in young children. The most commonly used behavioral tests relevant to ASD or schizophrenia in the mouse are performed in young adults. Yet it is worth noting that terminal axon branching defects can be detected as early as P21 in NUAK1^+/−^ mice, suggesting that some phenotypic alterations might be present before or around weaning. Importantly we also observed that the branching deficit persisted or even increased up to stage P90, corresponding to the age at which we assessed mouse behavior.

Many neurodevelopmental disorders have a strong sex bias. For instance ASD affects more frequently young males than females ^48^. Furthermore, schizophrenia displays some subtle differences in its incidence and clinical manifestations between male and female patients ^49^. The majority of the alterations we identified in NUAK1^+/−^ mice were similar in males and females, for example the reduction in terminal axon branching. However growth retardation was especially marked in NUAK1^+/−^ males, as was the increased startle response and decreased PPI reflex. The relevance of these findings for human pathology is currently unknown.

Gene-network analyses of genes linked to ASD or schizophrenia identified clusters of candidate genes involved in synapse formation, protein synthesis and RNA translation, signaling pathways, gene regulation and chromatin remodeling ^50–53^. This evidence points toward synaptic function and plasticity as key processes in the physiopathology of neurodevelopmental disorders. In line with that observation, a significant part of the clinical manifestation can be reversed after the development period, suggesting that synaptic plasticity plays an important role in the disease ^54^. Yet in parallel there is ample evidence that ASD and schizophrenia have a strong developmental component and that clinical manifestations may arise from early insults to neural circuits formation. The main consequence of NUAK1 heterozygosity is a reduction in terminal cortical axon branching, suggesting a direct link between axonal development and neurodevelopmental manifestations. Interestingly a similar reduction in terminal axon branching has been recently observed in a mouse model for the 22q11.2 deletion, strongly associated to schizophrenia ^55^. Our work strengthens the link between a disruption of cortical connectivity and disruption in cognitive and social behavior in the mouse and warrants further studies of axon development in the pathophysiology of neurodevelopmental disorders.

A recent study showed that NUAK1 mediates the phosphorylation of the microtubule associated protein TAU on S356 and might be involved in tauopathies such as Alzheimer’s disease (Lasagna-Reeves et al., 2016). These results suggest that NUAK1 over-activation might lead to neurodegeneration. In the present study, we have only examined the role of NUAK1 loss-of-function during embryonic and early postnatal development, but since its expression is maintained in the adult brain, future experiments should address NUAK1 function in adult circuits maintenance.

As a whole, we gathered evidence that expression of the autism-linked *Nuak1* Q433-Stop mutation disrupts cortical axon branching *in vivo*. Coupled with our observation that *Nuak1* heterozygosity impairs brain development, cortical axon branching *in vivo*, and that NUAK1^+/−^ mice have cognitive and social novelty defects strongly argues that *Nuak1* loss-of-function mutations could be causally linked to neurodevelopmental disorders in humans and support targeted efforts to identify additional genetic alterations in the *Nuak1* gene. Future experiments will need to determine if other rare *de novo* loss-of-function mutations identified in genes such as *Nuak1* are also haploinsufficient with regard to development of brain connectivity and if this is a general genetic mechanism underlying a fraction of neurodevelopmental and/or neuropsychiatric disorders.

## MATERIALS AND METHODS

### Animals

Mice breeding and handling was performed according to protocols approved by the Institutional Animal Care and Use Committee at The Scripps Research Institute, Columbia University in New York and the Ethics committee of the University of Lyon, and in accordance with National Institutes of Health guidelines and the French and European legislation. Time-pregnant females were maintained in a 12hr light/dark cycle and obtained by overnight breeding with males of the same strain. Noon following breeding was considered as E0.5. NUAK1^+/−^ mice (Nuak1^tm1Sia^) were obtained from the Riken Institute ^20^. Floxed NUAK1 mice were generated from targeted ES cells (Nuak1^tm1a(KOMP)Wtsi^) obtained from the KOMP Repository from UC Davis (https://www.komp.org/index.php, project ID CSD23401). Animals were maintained on a C57Bl/6J background.

### Image Acquisition and Analyses

Confocal images were acquired in 1024×1024 mode with a Nikon Ti-E microscope equipped with the C2 (Supplementary Fig 2 and Fig 7) or the A1R (all others) laser scanning confocal microscope using the Nikon software NIS-Elements (Nikon). We used the following objective lenses (Nikon): 10× PlanApo; NA 0.45, 20× PlanApo VC; NA 0.75, 60× Apo TIRF; NA 1.49. Time-lapse images were acquired in 512×512 mode with a Nikon Ti-E microscope equipped with an EM-CCD Andor iXon3 897 Camera. Mitochondria were imaged at 0.1 frames per second for 30 min. Analysis and tracking of confocal and time-lapse images was performed with NIS-Elements software. Representative neurons were isolated from the rest of the image using ImageJ. Contrast was enhanced and background (autofluorescence of non transfected neurons in culture) removed for better illustration of axons morphology. Kymographs were created with NIS-Elements.

### Behavior analyses

Two groups of age-related NUAK1 HET mice and WT littermates were submitted to behavioral assays in the Mouse Behavioral Assessment Core at The Scripps Research Institute in San Diego, CA. All quantifications were performed by experienced technicians blind to genotype. Mice were between age 10 and 12 weeks at the start of testing. Procedure, quantification method, order of experiments and group constitution is detailed in the Extended Material and Methods.

### DNA, Plasmids and cloning

The empty vector pCAG-IRES-GFP (pCIG2), CRE expressing vector pCIG2-CRE ^56^, mVENUS expressing vector pSCV2 ^57^, and mitochondria-tagged DsRed expressing pCAG-mitoDsRED ^13^ were described previously. peGFP-Flag-mNUAK1 vector was created by amplifying mouse NUAK1 cDNA by PCR and cloning into a peGFP-C1 vector between XhoI and EcoRI sites. A Flag-Tag was added at the N-terminal extremity of the protein by PCR. Kinase-dead (K85A), phosphorylation-defective (T212A) and STOP (Q434STOP) mutants were obtained by site-directed mutagenesis using the Quikchange II site-directed mutagenesis kit from Stratagene using the following primers (forward):

*NUAK1 K85A:* GCCGAGTGGTTGCTATAGCATCCATCCGTAAGGAC

*NUAK1 T212A:* GAAGGACAAGTTCTTGCAAGCATTTTGTGGGAGCCCAC

*NUAK1 Q434STOP:* CCCCTTTCAAAATGGAGTAAGATTTGTGCCGGACTGC.

Neuronal expressing vectors for NUAK1 (pCIG2-Flag-mNUAK1) were obtained by inserting wild-type and mutant NUAK1 cDNA into a pCIG2 vector between XhoI and EcoRI sites.

### Western blotting

Western blotting was carried out as previously published^58^. Cells and tissues were lysed in ice-cold lysis buffer containing 25 mM Tris (pH7.5), 2 mM MgCl2, 600 mM NaCl, 2 mM EDTA, 0.5% Nonidet P-40, and 1× protease and phosphatase mixture inhibitors (Roche). 20 µg aliquots of lysate were separated by electrophoresis on a 10% SDS-polyacrylamide gel and transferred on a polyvinylidene difluoride (PVDF) membrane (Amersham). Incubation with primary antibody was performed overnight in 5% milk-TBS-Tween. The next day, membranes were incubated at room temperature for 1 hour with HRP-linked secondary antibodies. Western-blot revelation was performed using ECL (Amersham). Imaging and analysis were performed on a chemidoc imager (Bio-rad) using the ImageQuant software.

### Ex Vivo Cortical Electroporation

Electroporation of dorsal telencephalic progenitors was performed as described previously ^59^. A mix containing 1 μg/μl endotoxin-free plasmid DNA (Midi-prep kit from Macherey-Nagel) plus 0.5% Fast Green (Sigma; 1:20 ratio) was microinjected in lateral ventricles of the brain of E15.5 embryos using the MicroInject-1000 (BTX) microinjector. Electroporations were performed on the whole head (skin and skull intact) with gold-coated electrodes (GenePads 5 × 7 mm BTX) using an ECM 830 electroporator (BTX) and the following parameters: Five 100 ms long pulses separated by 100 ms long intervals at 20 V. Immediately after electroporation, the brain was extracted and prepared as stated in the neuronal culture section below.

### Primary neuronal culture

Cortices from E15.5 mouse embryos were dissected in Hank's Buffered Salt Solution (HBSS) supplemented with Hepes (pH 7.4; 2.5 mM), CaCl_2_ (1 mM, Sigma), MgSO_4_ (1 mM, Sigma), NaHCO_3_ (4mM, Sigma) and D-glucose (30 mM, Sigma), hereafter referred to as cHBSS. Cortices were dissociated in cHBSS containing papain (Worthington) and DNAse I (100 μg/ml, Sigma) for 20 min at 37°C, washed three times and manually triturated in cHBSS supplemented with DNAse. Cells were then plated at 12.5 × 10^4^ cells per 35mm glass bottom dish (Matek) coated with poly-D-lysine (1 mg/ml, Sigma) and cultured for 5 days in Neurobasal medium supplemented with B27 (1X), N2 (1×), L-glutamine (2 mM) and penicillin (5 units/ml)-streptomycin (50 mg/ml). To transfect cultured neurons, we performed magnetofection using NeuroMag (OZ Bioscience) according to the manufacturer’s protocol.

### *In utero* cortical electroporation

Timed pregnant NUAK1^+/−^ or NUAK1^F/F^ females were used for In utero cortical electroporation. A mix containing 1 µg/µl endotoxin-free plasmid DNA (Midi-prep kit from Macherey-Nagel) plus 0.5% Fast Green (Sigma; 1:20 ratio) was injected into one lateral hemisphere of E15.5 embryos using a picospritzer. Electroporation was performed using an ECM 830 electroporator (BTX) with gold-coated electrodes (GenePads 3 × 5 mm BTX) to target cortical progenitors by placing the anode (positively charged electrode) on the side of DNA injection and the cathode on the other side of the head. Four pulses of 45 V for 50 ms with 500 ms interval were used for electroporation. Animals were sacrificed 21 or 90 days after birth by terminal perfusion of 4% paraformaldehyde (PFA, Electron Microscopy Sciences) followed by 2 hours post-fixation in 4% PFA.

### Statistical Analyses

Sample size is detailed in the figures. Statistical tests were performed using Prism (GraphPad Software)̤ ns: p>0.05, *: p<0.05, **: p<0.01, ***: p<0.001. All statistical tests are two-tailed. Normality test (D’Agostino & Pierson) was performed prior to analysis to discriminate between parametric or non-parametric assay. Retrospective comparison of variance was performed with a F-test.

## ACKNOWLEDGEMENTS

The authors express their gratitude to members of the Polleux and Courchet labs for useful comments and discussion. We are grateful to J. Gogos and A. Diamantopoulou for critical reading of the manuscript. This work was supported by NIH-R01NS089456 (FP), NIH-F32NS080464 (TL), NIH-K99NS091526 (TL), the Fondation pour la Recherche Médicale (AJE20141031276) and ERC Starting Grant (678302-NEUROMET). J.C. was the recipient of a grant from the Philippe Foundation. VC, AR, TL, FP and JC conceived the experiments and interpreted the results. VC, TL, and JC performed the experiments. AR, VC and PDC performed behavior analyses. JC and FP prepared the manuscript.

